# The interplay between lumen pressure and cell proliferation determines organoid morphology in a multicellular phase field model

**DOI:** 10.1101/2023.08.17.553655

**Authors:** Sakurako Tanida, Kana Fuji, Linjie Lu, Tristan Guyomar, Byung Ho Lee, Alf Honigmann, Anne Grapin-Botton, Daniel Riveline, Tetsuya Hiraiwa, Makiko Nonomura, Masaki Sano

**Author notes:** (TH); (MN); (MS).

## Abstract

Organoids are ideal systems to predict the phenotypes of organs. However, there is currently a lack of understanding regarding the generalized rules that enable use of simple cellular principles to make morphological predictions of entire organoids. Therefore, we employed a phase field model with the following basic components: the minimum conditions for the timing and volume of cell division, lumen nucleation rules, and lumenal pressure. Through our model, we could compute and generate a myriad of organoid phenotypes observed till date. We propose morphological indices necessary to characterize the shapes and construct phase diagrams and show their dependencies on proliferation time and lumen pressure. Additionally, we introduced the lumen-index parameter, which helped in examining the criteria to maintain organoids as spherical structures comprising a single layer of cells and enclosing an intact lumen. Finally, we predict a star-like organoid phenotype that did not undergo differentiation, suggesting that the volume constraint during cell division may determine the final phenotype. In summary, our approach provides researchers with guidelines to test the mechanisms of self-organization and predict the shape of organoid.

**Author summary:** In nature, a wide variety of organ morphologies are observed. Owing to the complexity of the process underlying the acquisition of organs’ morphology, it is challenging to investigate the mechanisms that lead to such variations. A promising approach to study these variations is the use of “computational organoid” study, which is the computational-based study of self-organizing shapes in multicellular assemblies and fluid-filled cavities called lumens that develop from a few proliferating cells. This study explores general mechanisms that dictate how various mechanical factors affect the growing self-organized multicellular assembly. We relied on computer simulations of the mathematical model called multicellular phase-field model with lumens and explored the mechanical factor effects, such as the lumen pressure while considering the time and volume conditions required for cell division. These simulations generated and categorized a wide range of organoid phenotypes based on the varying lumen pressure and cell division conditions. These phenotypes were characterized into seven distinct classes, based on the morphological index sets, including a cellular monolayer/multilayer surrounding single or multiple lumens and branch formation. These phenotypes were obtained without the assumption of differentiation. Our study elucidates the mechanisms underlying the organoid and organ formation with different shapes, thereby highlighting the significance of mechanical forces in shaping these complex biological structures.

## Introduction

Morphology plays a crucial role in organ function and there is a wide variety of organ morphologies in nature. As the processes involved in the acquisition of real organs’ morphologies are extremely complex, the underlying mechanisms can be elucidated through the exploration of simplified systems such as organoids. Organoids are generally defined as an *in-vitro* cell assembly with a specific configuration that develops from a few stem/progenitor cells through self-organization, and could offer a new understanding regarding this [1, 2]. As shown in Fig. 1, organoids exhibit a variety of morphologies and are therefore considered a potential model to study the principles underlying the determination of various tissue morphologies. While simple examples of these morphologies include cell aggregates such as tumor spheroids [3], many organoids have lumens, which are liquid-filled cavities surrounded by tissue cells. Simple monolayer lumens are found throughout the body in organs such as the thyroid gland [4–6] and often form *in vitro* from cells that would naturally form tubes *in vivo*.

**Fig 1.**
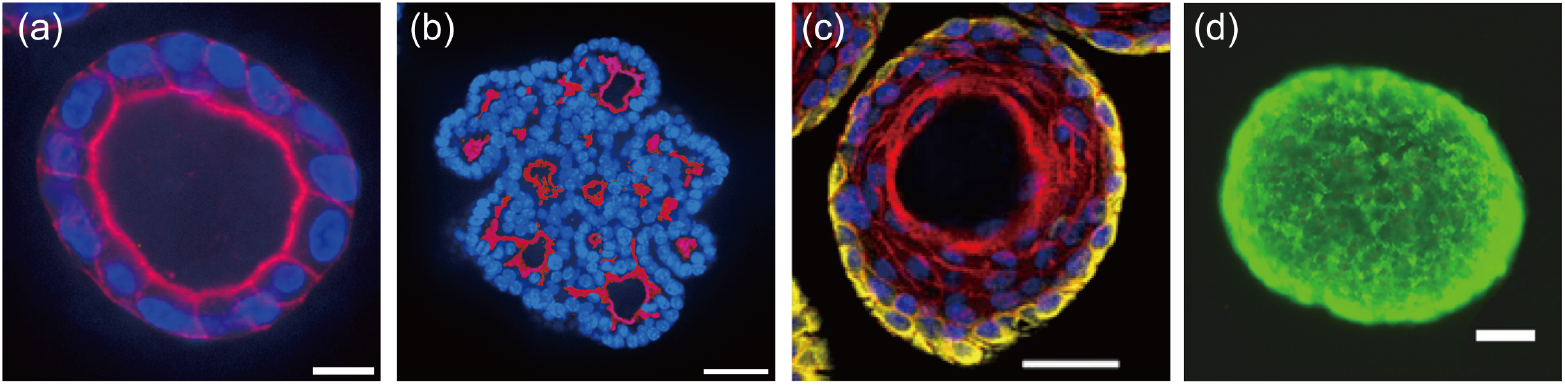
Variety of morphologies observed in *in vitro* organoids. Cross-sectional view of different organoids with the lumen surface labeled in red. (a) Snapshot of an Madin-Darby canine kidney (MDCK) cyst with a single lumen labeled with F-actin (red) and DAPI (blue). Scale bar: 10 *μ*m. (b) Snapshot of a pancreatic organoid with multiple lumens labeled with Ezrin (red) and Hoescht (blue). Scale bar: 40 *μ*m. (c) Snapshot of a murine epidermal organoid with a single lumen surrounded by multiple cell layers labeled with F-actin (red), DAPI (blue), and Keratin-5 (yellow). Adapted with permission from Boonekamp KE et al. [13]. Scale bar: 50 *μ*m. (d) Human breast adenocarcinoma tumor spheroid with metabolically viable cells shown in green and the central cells of the spheroid remained viable. Adapted with permission from Gong X et al. [3]. Scale bar: 100 *μ*m.

The isotropic environment of suspension or gel cultures simplifies their geometry from a tube to a sphere. Intestinal organoids retain some of their folding structure from the intestinal crypts and villi and exhibit short tubular structures bulging from a cyst [7–9]. Small buds growing on a cyst structure are also found in organoids of the stomach and liver organoids [10–12]. More complex structures may also form *in vitro*. For example, epidermal organoids exhibit a central cyst surrounded by multiple cell layers [13]. Others can have multiple lumens, or even a network of lumenal structures.

Multi-lumenal structures within a multilayered stratified epithelium are observed in the pancreas during development *in vivo* and in pancreatic organoids *in vitro* [14, 15]. A network of lumenal structures are observed during angiogenesis [16, 17], and in the formation of the lung [18–20] and pancreas both *in vivo* and *in vitro* [14, 15, 21, 22].

While the mechanisms of lumen formation have been explored in great depth notably using the Madin-Darby canine kidney (MDCK) lumen system, the mechanisms that control the diversity of lumen shapes has not been comprehensively studied [23–25]. The diversity of organoid models enables the study of this *in vitro*, which makes the topic timely, notably considering the tissue mechanics.

A hallmark of organoids is the proliferation of their cells accompanied by morphogenesis and these structures are formed by a variety of factors. Mechanics and hydraulics play pivotal roles in the growth of cell assemblies, including organs and organoids. Moreover, some organs/organoids can sense the mechanical environment and applied forces and thus change their morphologies accordingly [8, 9, 26–31]. When starting with epithelial cells that can polarize and form a lumen, micro-lumens are initially created within or between the cells and then expand and form a physiologically functioning structure [32]. This expansion is driven by the osmotic pressure differences generated through ion transport mechanisms on the cell surface that causes water to flow into the lumen from the surrounding environment [32–35]. In a system such as this, the balance of forces, such as that between the lumen pressure and tissue elasticity of the cell monolayer, plays an important role in shaping the entire system. Additionally, the balance of kinetics must also be considered; for example the speed of lumen expansion must match that of the cell proliferation rate for the stable existence of the lumen; in the absence of this balance, lumen could disappear or leak out from between the cell junctions [36]. These factors can be considered independently of the cell state/identity, whether the cell is a stem cell or has differentiated from pluripotency. Therefore, in this study, we do not focus on the cell state but rather consider the underlying mechanistic factors. It is important to note that this assumption does not limit the scope of the study regarding the cell states in which differentiation does not occur. Rather, it is a framework that enables the discussion of the topic regardless of the cell state.

To reveal the mechanisms that can determine the morphology of the organoids or real organs observed in each experiment, careful theoretical discussions are necessary to integrate these factors. However, such discussions are currently unavailable. Specifically, the following elements are missing: (i) fundamental knowledge on what can mechanically emerge in the system that comprises of proliferating cells under geometrical constraints and (ii) a systematic manner to characterize the observed morphology.

In this study, we addressed this issue through the analysis of a computational model of organoids, *i*.*e*., growing computationally simulated cell assemblies. Assuming a typical organoid culture environment, we set a few cells (four cells in this study) as the initial condition and computationally simulated the self-organized multicellular structures after multiple rounds of cell proliferation. Simulations were performed by applying the computational framework called multicellular phase-field model, which can simulate the shape of each cell and lumen using the continuum field in space [37, 38]. Although there are other models simulate the applicable assembly structure [39–43] and lumen-containing organoids [44], such as the cellular vertex and Potts models, considering the purpose of this study to determine the effect of mechanical factors including pressure, we adopted the phase field model, which can model the pressure of both the cells and lumens [43]. The cellular vertex model can clearly describe the forces generated by the cytoskeleton and applied to the vertex, but the shape of cells and lumens are limited to polygonal shapes. The cellular Potts model allows for the easy introduction of rules for cell–cell and cell–lumen interactions, but it is challenging to clearly describe the balance of the forces. We observed that the growing cell assembly self-organized into varieties of morphologies only through the coupling between the mechanics/kinetics of growth and geometrical constraints by the multicellular phase field model. These morphologies were characterized by the proposed indices. These results may provide a manner to study the mechanics/kinetics-based principles that determine the formation process and final forms of the organoids, and be applied to gain a deeper understanding of the morphogenetic mechanisms of functional organs.

In the following sections, we discuss the computational simulations. We present the results and analysis in the Results and Discussion sections, respectively. This includes the phase diagram, morphology, and indices used to characterize the model. We also explore the effect of introducing noise to the cell division process, as well as the mechanism that maintain the monolayer morphology. In the Discussion section, we further address the challenge of characterizing the organoid morphologies under the experimental conditions. We also emphasize the importance of considering the model limitations and investigating the role of the extracellular matrix. These insights provide valuable guidance for future research endeavors. In the last, we comprehensively describe the mathematical model in the Materials and Methods section.

## Results

We began with four cells at the center of the simulation domain in a two-dimensional system. Each cell adhered to two neighboring cells, forming a circular shape that was divided into four equal quadrants. The space and time step sizes, d*x* and d*t*, were set to 0.02 arb. unit of length and 0.01 arb. unit of time, respectively. The simulation domain was selected to be sufficiently large compared to that of a single cell, and the simulation boundary was square, with a size of 40 × 40. When the organoid reached the simulation boundary, the simulation was set to stop. For the analysis of the volume and perimeters of the cells and lumen, we defined the region of a cell and a lumen, where *u*_*i*_ *>* 0.2 was satisfied. For convenience, we refer to the area in two dimensions as the ‘volume’ in this paper.

### Patterns of organoids

We numerically simulated our organoid model by varying the parameters of *t*_*d*_ and *ξ* from 0 to 300 arb. unit of time and from 0.27-0.40 arb. unit of pressure, respectively, with increments of 20 and 0.01. Figure 2 shows the phase diagram and typical pattern of each organoid morphology. Based on the morphology features, we classified the simulated organoid patterns into seven distinct types: star-shaped, monolayer lumen, branched multi-lumen, multilayer multi-lumen, multilayer no-stable-lumen, multilayer single-stable-lumen, and ruptured sheet with expanding lumen. The ruptured sheet with expanding lumen was excluded from our analysis because no organoid growth is typically observed after rupture. The star-shape and monolayer cyst organoids exhibited a single lumen surrounded by a single cell layer, with the former breaking rotational symmetry and the latter having a round shape [Fig. 2(b,c)]. Both branched multi-lumen and multilayer multi-lumen organoids had multiple lumens within the cell layers, with the former increasing the lumen area over time and the latter decreasing the lumen area over time [Fig. 2(d,e)]. The multilayer no-stable-lumen corresponded to a state in which the lumen area eventually disappeared and multiple cell layers were persistently present, as the name implies [Fig. 2(f)]. Similarly, the multilayer single-stable-lumen was a state in which one lumen area was surrounded by multiple cell layers [Fig. 2(g)]. This phase diagram highlights the importance of the cell proliferation rate and lumen growth rate to determine the morphology of organoids. To further explore the mechanism of the formation of different shapes, we used specific indices to characterize each morphology, which are discussed in the following subsections.

**Fig 2.**
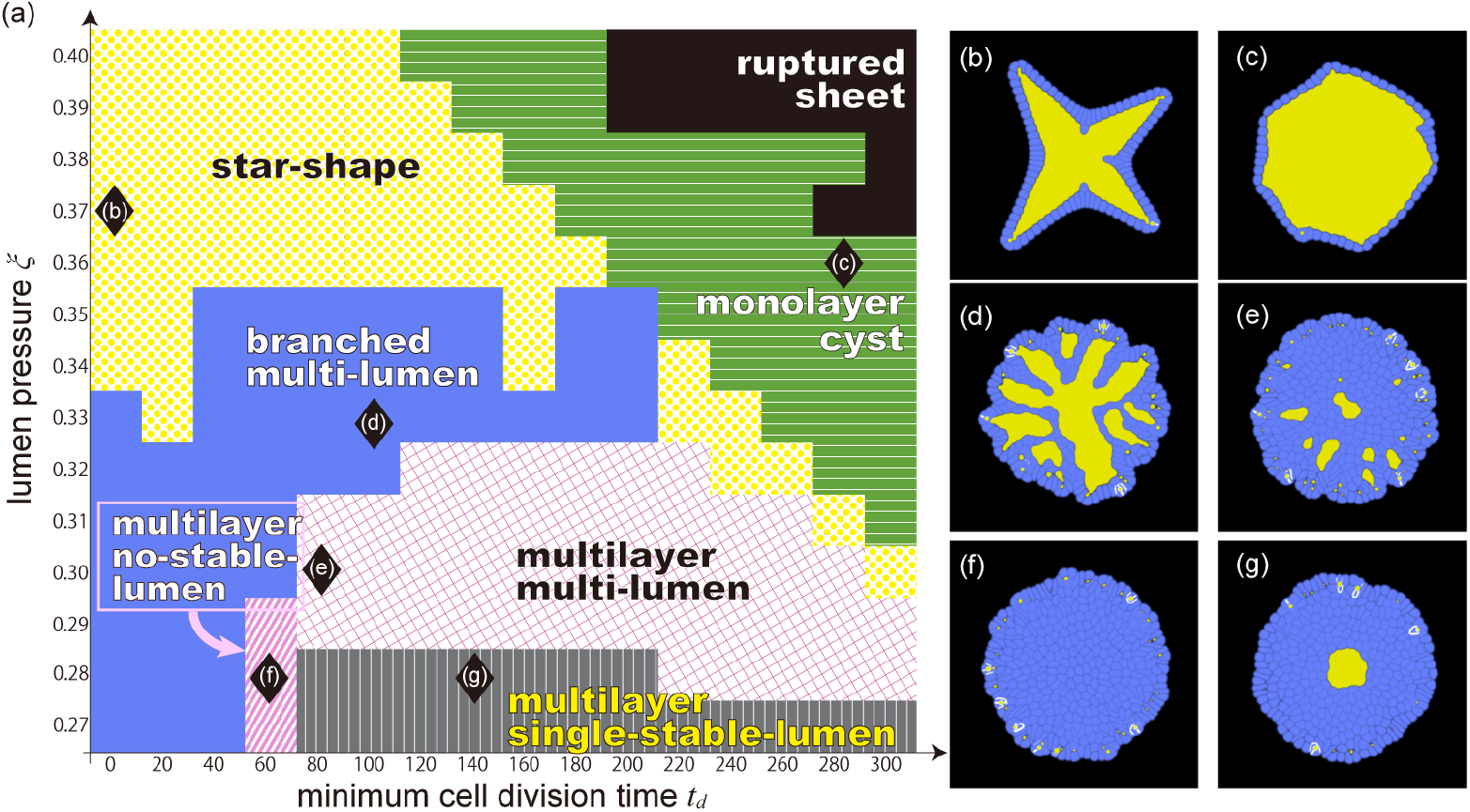
Overview of the morphologies produced by this model. (a) Phase diagram of the organoid morphology and typical pattern of each phase. Each domain corresponds to; star shape (yellow dots), monolayer cyst (green horizontal stripe), branched multi-lumen (red solid), multilayer multi-lumen (magenta lattice), multilayer no-stable-lumen (blue tilted stripe), and multilayer single-stable-lumen (navy vertical stripe). The black and white star markers correspond to the parameter sets where the organoids of (b)-(g) emerge. The final states of (b) star shape organoid formed when (*ξ, t*_*d*_) = (0.37, 0), (c) Monolayer cyst organoid formed when (*ξ, t*_*d*_) = (0.36, 280), (d) branched multilayer organoid formed when (*ξ, t*_*d*_) = (0.33, 100), (e) multilayer multi-lumen organoid formed when (*ξ, t*_*d*_) = (0.30, 80), (f) multilayer no-stable-lumen organoid formed when (*ξ, t*_*d*_) = (0.28, 60), (g) multilayer single-stable-lumen organoid formed when (*ξ, t*_*d*_) = (0.28, 140). The blue and yellow regions in (b-g) represent the cells and lumen, respectively.

Figure 3 illustrates the time evolution of each morphology. In the beginning, four initial cells were clustered in the center of the computational area. In most cases, the first micro-lumen was created when the first four cells divided and merged in the center, surrounded by more cells that divided radially to create a monolayer. This process occurred in the star-shape, monolayer cyst, branched multi-lumen, and multilayer single-stable-lumen morphologies [Fig. 3(a), (b), (c), and (f)]. After the initial formation of micro-lumens, the star-shaped organoid branched outward, while the single-layer cyst grew in a relatively stable shape. In the branched multi-lumen morphology, multiple lumens were generated in the outer cell layers, some of which merged with the center lumen. In the multilayer single-stable-lumen, subsequent micro-lumens that were generated in the process were not stable and disappeared without growing. In contrast, in the multilayer multi-lumen and multilayer no-stable lumen morphologies [Fig. 3(d) and (e)], a lumen did not initially form in the center due to the pressure from the surrounding cells. Instead, stable lumens formed in the outer cell layers during the subsequent phase for the multilayer multi-lumen morphology.

**Fig 3.**
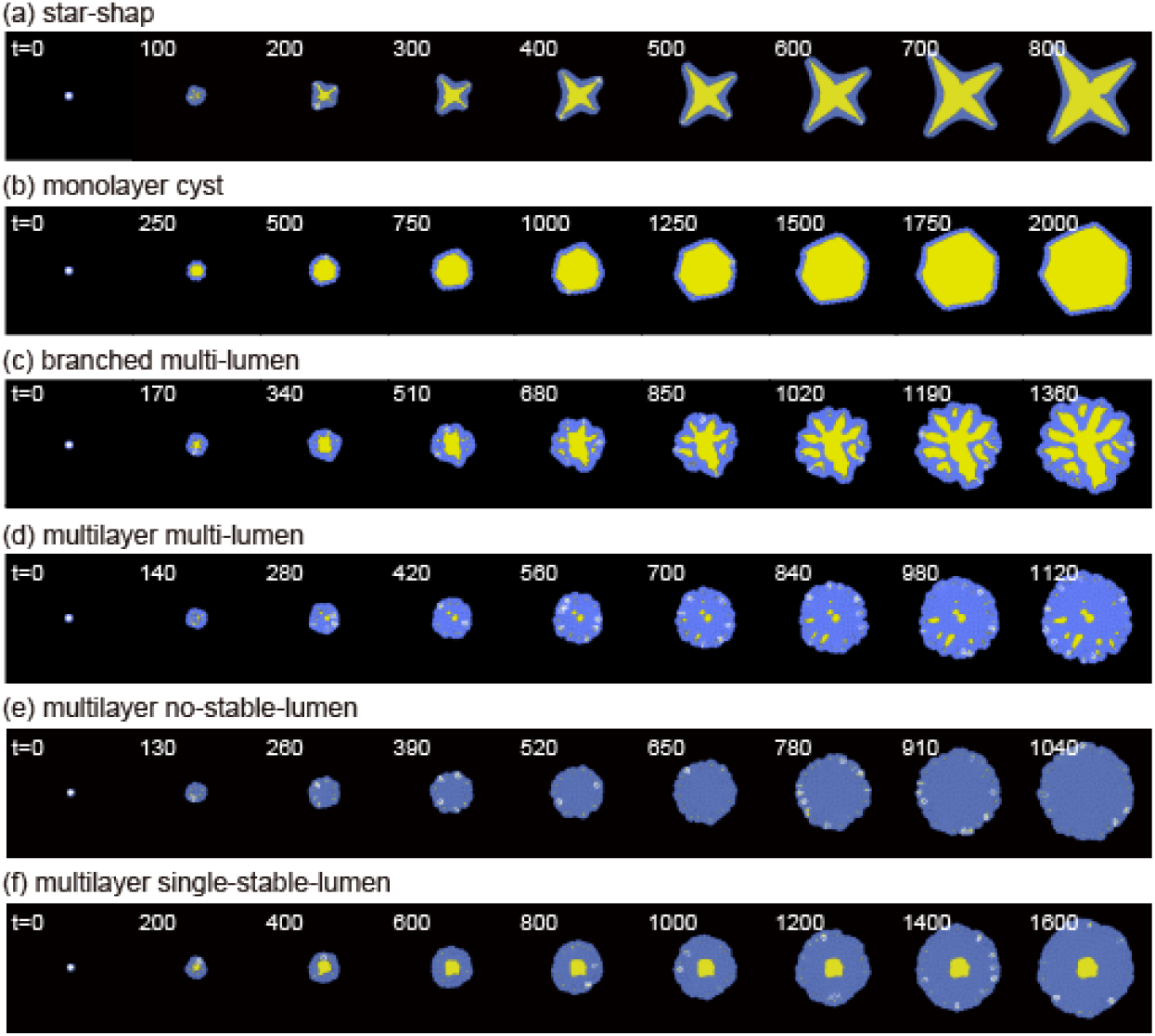
Time evolution of the organoid growth process. The blue and yellow regions in (a-g) represent the cells and lumen, respectively. (a) star-shape (*ξ, t*_*d*_) = (0.37, 0), (b) monolayer cyst (*ξ, t*_*d*_) = (0.36, 280), (c) branched multi-lumen (*ξ, t*_*d*_) = (0.33, 100), (d) multilayer multi-lumen (*ξ, t*_*d*_) = (0.30, 80), (e) multilayer no-stable-lumen (*ξ, t*_*d*_) = (0.28, 60), and (f) multilayer single-stable-lumen (*ξ, t*_*d*_) = (0.28, 140). See supplemental movies 1-6.

### Indices to characterize organoid morphology

The patterns that appeared in the simulations were categorized based on the extent of lumen occupancy, number, and sphericity. Table 1 provides a summary of the range of each index for each organoid shape.

**Table 1.**
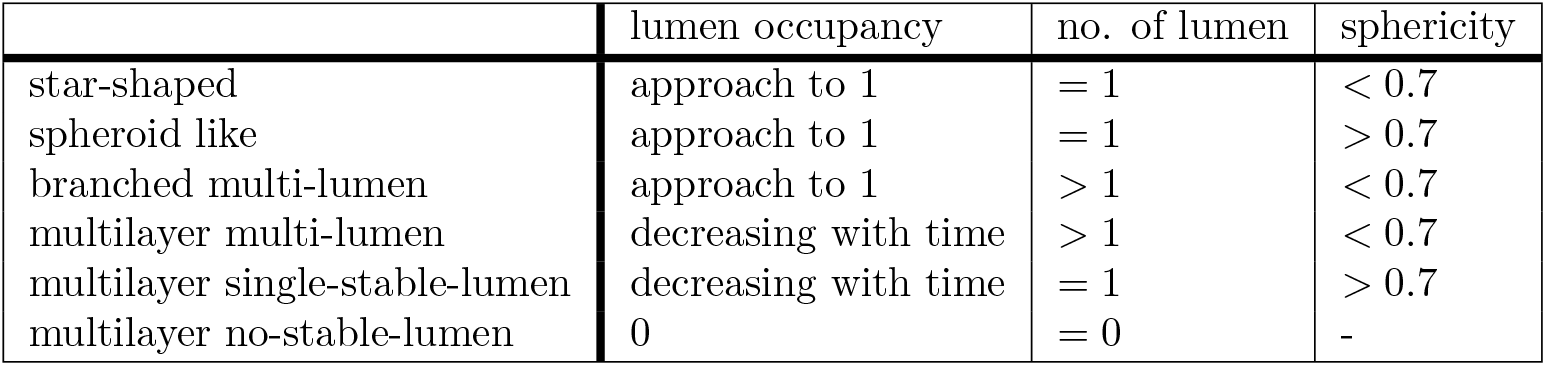
Summary of the behaviors of the indices to characterize the organoid morphology.

### Lumen Occupancy

The first index used to classify the morphologies is the lumen occupancy, which is defined as the volume of lumen over the volume of the organoid in 2D, and is expressed as:

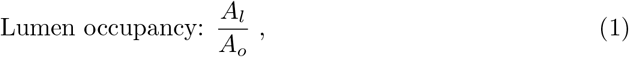

where *A*_*l*_ and *A*_*o*_ are the areas of the lumen and organoid, respectively [Fig. 4(a)]. Here, *A*_*o*_ includes the area of *A*_*l*_. The simulated organoids are growing systems with no steady states. Thus, we focused on the time variation to capture the growing morphology.

**Fig 4.**
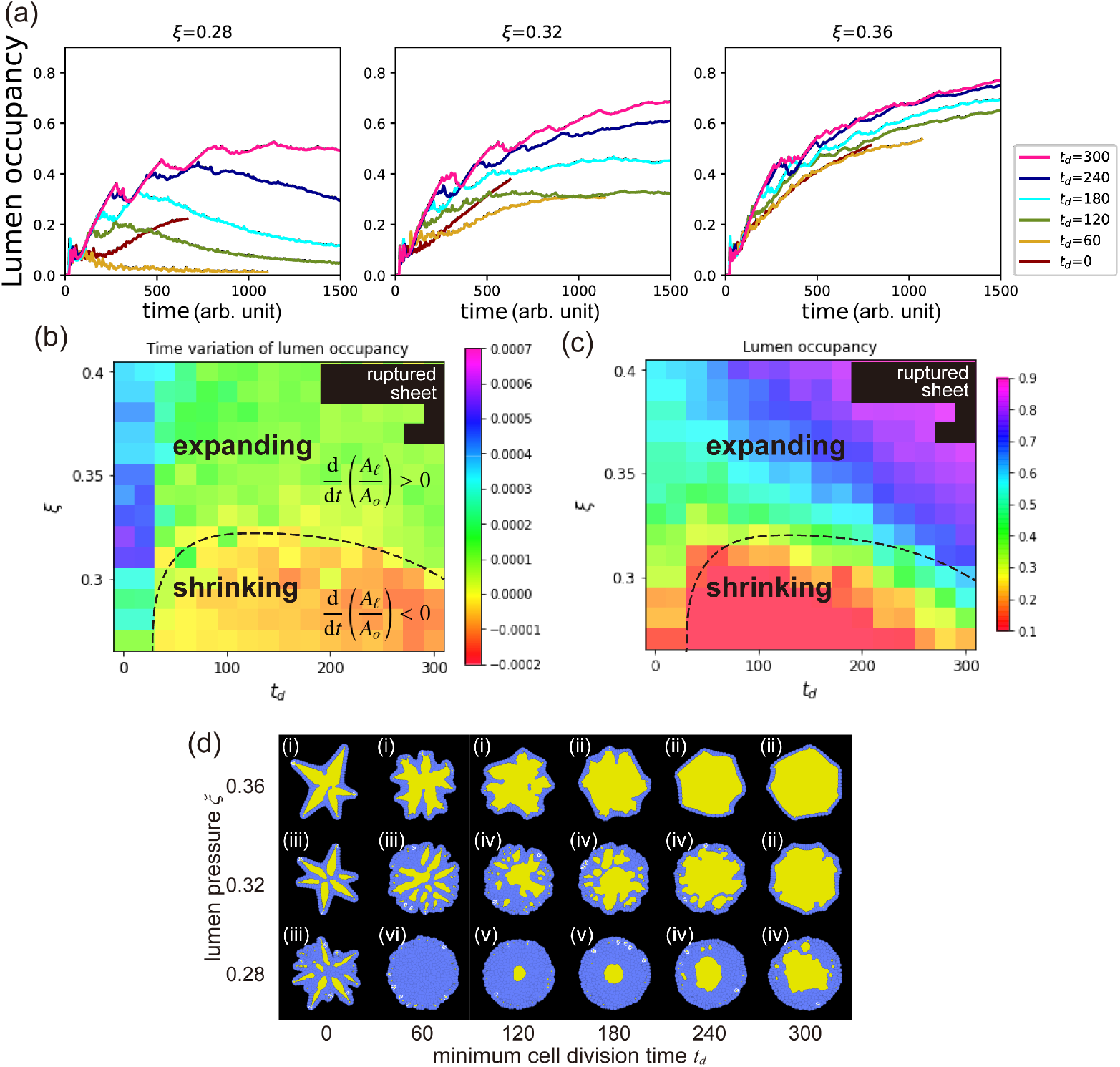
Lumen occupancy. (a) Time evolution of the lumen occupancy at the various *t*_*d*_ and *ξ*. (b) Time variation rate of lumen occupancy for the various values of the minimum cell division time *t*_*d*_ and lumen pressure *ξ*. Black dashed line represents the contour line of zero. The lumen occupancy increases and decreases with time to above and below the black dashed line, respectively. (c) Value of the lumen occupancy at the final state for the various *t*_*d*_ and *ξ*. (d) Snapshot of each parameter set of (c). The number labels represent the class of the organoid morphology: (i) star-shape, (ii) monolayer cyst, (iii) branched multi-lumen, (iv) multilayer multi-lumen, (v) multilayer single-stable-lumen, and (vi) multilayer no-stable-lumen.

Figure 4(c) shows how the lumen occupancy changes over time for the different lumen pressures (*ξ*) and minimum time for cell division (*t*_*d*_). At *ξ* = 0.28, the lumen occupancy increased in the beginning, but decreased in the subsequent phase, except for *t*_*d*_ = 0. Organoids at *ξ* = 0.28 exhibited multiple cell layers (lower row in Fig.4(e)). In most cases at *ξ* = 0.28, the lumen occupancy initially increased but then decreased in the subsequent phase except for *t*_*d*_ = 0. The simulation at (*ξ, t*_*d*_) = (0.28, 0) was terminated before lumen occupancy could start to decrease because the organoid grew larger than the simulation area. For *ξ* = 0.32, the lumen occupancy increased in the beginning, and the slope gradually decreased before reaching a constant value. Organoids at *ξ* = 0.32 and *t*_*d*_ values of 0, 60, 120, 180, and 240 harbored multiple cell layers, while those at *t*_*d*_ = 300 had a single cell layer [middle row in Fig. 4(e)]. When *ξ* = 0.36, the lumen occupancy increased over time, regardless of *t*_*d*_, which resulted in monolayer organoids [upper row in Fig. 4(e)].

To better understand the behavior of the organoids in the subsequent phases, we analyzed the rate of change in the lumen occupancy during the last 200 arb. units of time. Where each unit corresponds to a time step interval of 0.01 in the arb. units of time used for time discretization in this study. The results are presented in a heat map in Fig. 4(b), where two main regions can be identified: one where the lumen occupancy increases (above the black dashed line), and the other where it decreases (below the black dashed line) as the lumen pressure decreases.

We also examined the final lumen occupancy for each of the parameter sets. Fig. 4(d) shows that lumen occupancy was generally higher at higher values of *ξ* and *t*_*d*_. As both parameters increase, lumen occupancy also tends to increase. The black dashed line represents the point where the lumen occupancy does not change over time.

By comparing the variation in the lumen occupancy, the growth of the organoids can be discussed. When the lumen occupancy expands over time, there is a greater chance of the cells being a monolayer. Conversely, when it decreases over time, the cell layer becomes thicker, indicating the presence of multiple cell layers in the organoid. This criterion can be applied to both three-dimensional simulations and experimental systems to classify the growing organoids. Measurements can be taken by determining the volume of the lumen and the entire organoid using three-dimensional scanning, or by measuring the area of the lumen and organoid in cross-section where the cells can be tracked.

### Number of lumens

The second index of our analysis is the number of stable lumens present, as shown in Figure 5(a). The stable lumens were defined in this study as lumens that have twice the volume as that of the micro-lumens generated immediately after cell division. We excluded micro-lumens from our lumen counts because they are not necessarily stable over a long period. Figure 5(b) shows the number of stable lumens in the final phase of each parameter set. In general, the number of lumens in the final state is either one at high and low lumen pressures or more than one at intermediate lumen pressure. However, the organoids at (*ξ, t*_*d*_) = (0.27, 60), (0.28,60), and (0.29,60) did not have any stable lumens present.

**Fig 5.**
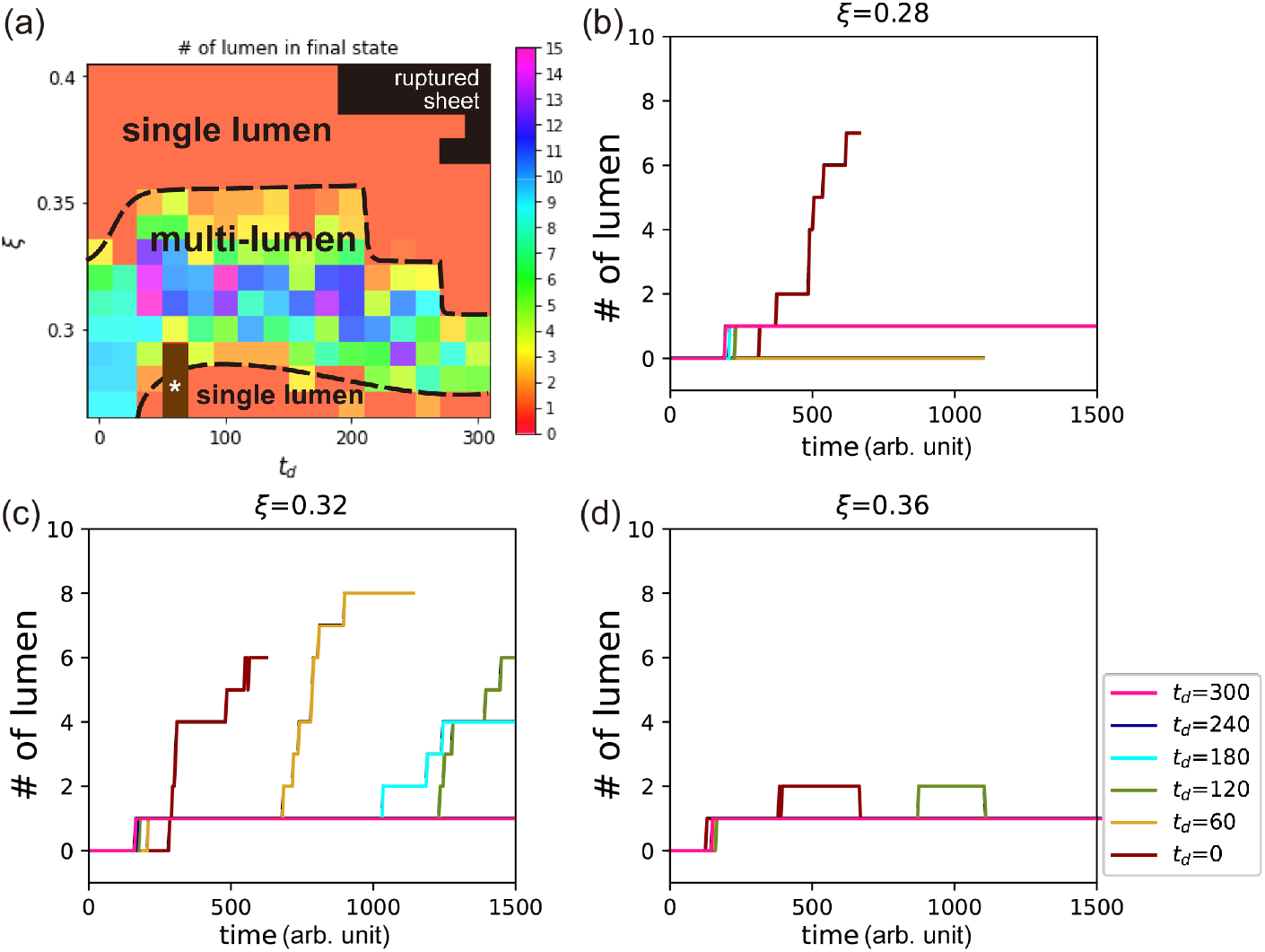
Number of lumens. (a) Number of lumens present at the various values of the minimum cell division time *t*_*d*_ and lumen pressure *ξ*. Brown region with a star marker, *, represents zero. (b-d) Time evolution of the number of lumen at the various *t*_*d*_ and *ξ*. See their morphology in Fig. 4(e).

Figure 5(c) displays the number of lumen over time at *ξ* = 0.28, 0.32, and 0.36. At *ξ* = 0.28, the organoids with a multilayer no-stable-lumen configuration are represented by the line at *t*_*d*_ = 60 in the left panel of Fig. 5(c), where the number of lumen remains zero for nearly the entire time. For *t*_*d*_ = 120 and 180 at *ξ* = 0.28, the number of lumens was consistently one. However, for *t*_*d*_ = 0, 240, and 300 at *ξ* = 0.28, the number of lumen tended to increase as time increased. In the center panel of Fig. 5(c), the number of lumen at *ξ* = 0.32 was mostly more than one, except at *t*_*d*_ = 300. In contrast, for organoids with a single lumen, such as those at higher values of *ξ*, such as (*ξ, t*_*d*_) = (0.36, 300), the number of lumens remained at one for nearly entire time, as shown in the right panel of Fig. 5(c).

The number of lumens is an important indicator of the complexity of the organoid’s inner structure, as it reflects whether the internal structure is composed of only cells, a cyst-like lumen, or more complex structures. Counting the number of lumens in a cross-section can therefore provide valuable insight into the developmental processes and morphogenesis of organoids in both three-dimensional simulations and experimental systems.

### Sphericity

The third index of our analysis is sphericity, which is defined as

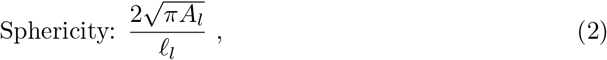

in 2D, where *ℓ*_*l*_ is the perimeter of the lumen. When an organoid has a single lumen that is circular in shape, its sphericity index is one [Fig. 6(a)]. However, if the organoid has a single lumen with a complex shape or multiple lumens, the sphericity index will be smaller than one. Note that the sphericity index cannot be defined for organoids with no lumens.

**Fig 6.**
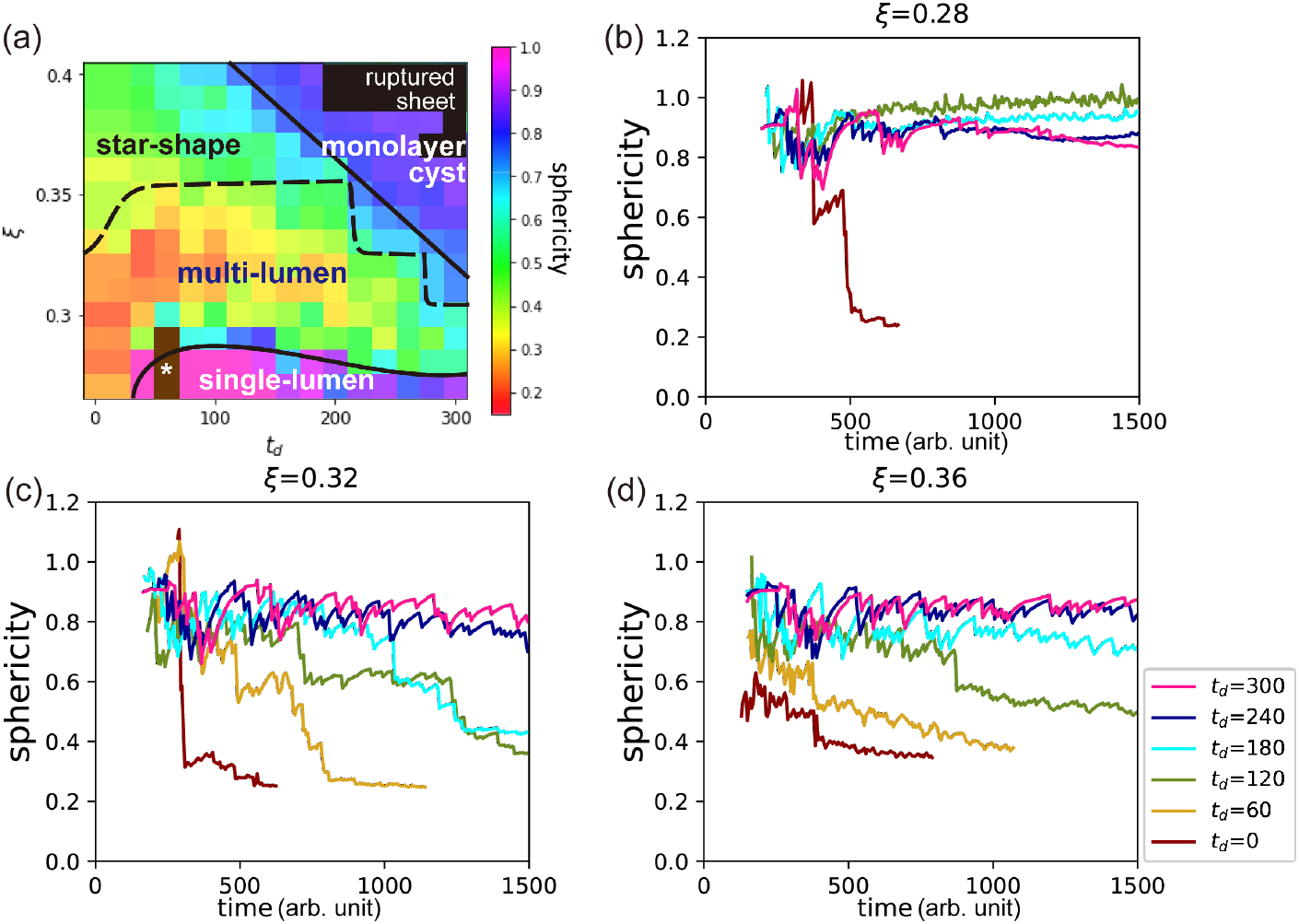
Sphericity. (a) Sphericity of organoids at various values of the minimum cell division time *t*_*d*_ and lumen pressure *ξ*. The brown region with a star marker, *, represents the zero lumen region. (b-d) Time evolution of the sphericity at various values of *t*_*d*_ and *ξ*. See their morphology in Fig. 4(e).

The sphericity index is crucial to distinguish between the star-shaped and monolayer lumen organoids. For lumen pressures higher than the black dashed line, which separates the multi-lumen from single-lumen configurations, the sphericity is generally higher at the lower values of *t*_*d*_ compared to the higher values of *t*_*d*_. The transition from star-shaped to monolayer single-lumen organoids occurs at a crossover point. At low lumen pressures, the sphericity separates the multilayer multi-lumen and multilayer single-stable-lumen configurations, as indicated by the lower black solid line in Fig. 6(b).

We define sphericity of the boundary between the star-shaped and monolayer single-lumen phases as 0.7, based on the jump in sphericity between the configuration of multilayer multi-lumen and multilayer single-stable-lumen. This boundary is shown by the upper black solid line in Fig. 6(b).

The sphericity index at (*ξ, t*_*d*_) = (0.36, 240), (0.36, 300), and (0.32, 300), where the monolayer single-lumen emerges, maintains a value of approximately 0.8 from the early simulation phase [Fig. 6(c)]. Conversely, at (*ξ, t*_*d*_) = (0.36, 120) and (0.36, 180), the sphericity decreases over time, indicating the transition from monolayer single-lumen to star-shape organoids. It also decreases with time in the pattern of multi-lumen organoid; for example, *ξ* = 0.32 and 0 ≤ *t*_*d*_ ≤ 240 [the center panel of Fig.6(c)]. At *ξ* = 0.28 and *t*_*d*_ *>* 60, where the multilayer single-stable-lumen organoids form, sphericity reaches 0.9 in the early phase and maintains this value throughout [left panel of Fig. 6(c)].

In this study, we found that the sphericity index was a crucial parameter to distinguish between the different configurations of organoids, such as between the star-shaped and monolayer lumen organoids. We defined the sphericity of the boundary between these phases based on the jump in sphericity between the different configurations. The sphericity index provides valuable insight into the structural complexity of the organoids and can help understand how the structure of the cell sheet affects their fate and the organoid functions. Therefore, examining the sphericity in three-dimensional simulations and experiments can be a useful tool to study organoid morphogenesis.

### Effect of noise added to the minimum volume condition

In previous simulations, the condition of the minimum volume required for cell division was constant and *V*_*d*_ = 2.9 arb. unit of volume for all cells although a small amount of noise was added to *t*_*d*_. However, we found that the condition of the minimum time required for cell division was more easily satisfied compared with that of the minimum cell volume condition for cell division for many of the classified patterns. Therefore, in those cases, the minimum volume condition dominates, and the resulting growth dynamics were almost deterministic. This is in contrast to real organoid systems, where high variability and heterogeneity exists within the same experimental conditions, which suggests the influence of fluctuations [45]. To investigate this, we introduced variability in the minimum volume condition for cell division, where *V*_*d*_ was assumed to follow a normal distribution with a mean of 2.85 and standard deviation of 0.025. In this section, we explored the impact of this variability on the shape of the organoids. For example, a pattern such as the star-like shape breaks rotational symmetry, and it is therefore possible that fluctuations in cell volume could contribute to the observed variability and heterogeneity in real organoid systems.

As shown in Fig. 7(a), the phase diagram exhibits the same morphology classes as discussed in the previous subsection, with no alteration in the relative positioning of the classes compared to Fig. 2. Each class appears at a higher lumen pressure region compared to Fig. 2 because of the reduced average threshold value and subsequent increase in the possibility of cell division. Nevertheless, the inclusion of noise in the minimum volume condition does not alter the qualitative characteristics of the system.

In contrast, there are notable differences in the morphological details of certain classes. The star-shaped organoids with noise tended to have more arms [Fig. 7(b)] than those without added noise [Fig. 2(b)]. Moreover, the direction of the extensions of the branches were not restricted to specific directions, which restored the rotational symmetry in a statistical sense. Furthermore, monolayer single-lumen organoids exhibited a rounder shape without facets, and the same is true for the branched multi-lumen and multilayer multi-lumen morphologies, where the facets were absent in each lumen.

**Fig 7.**
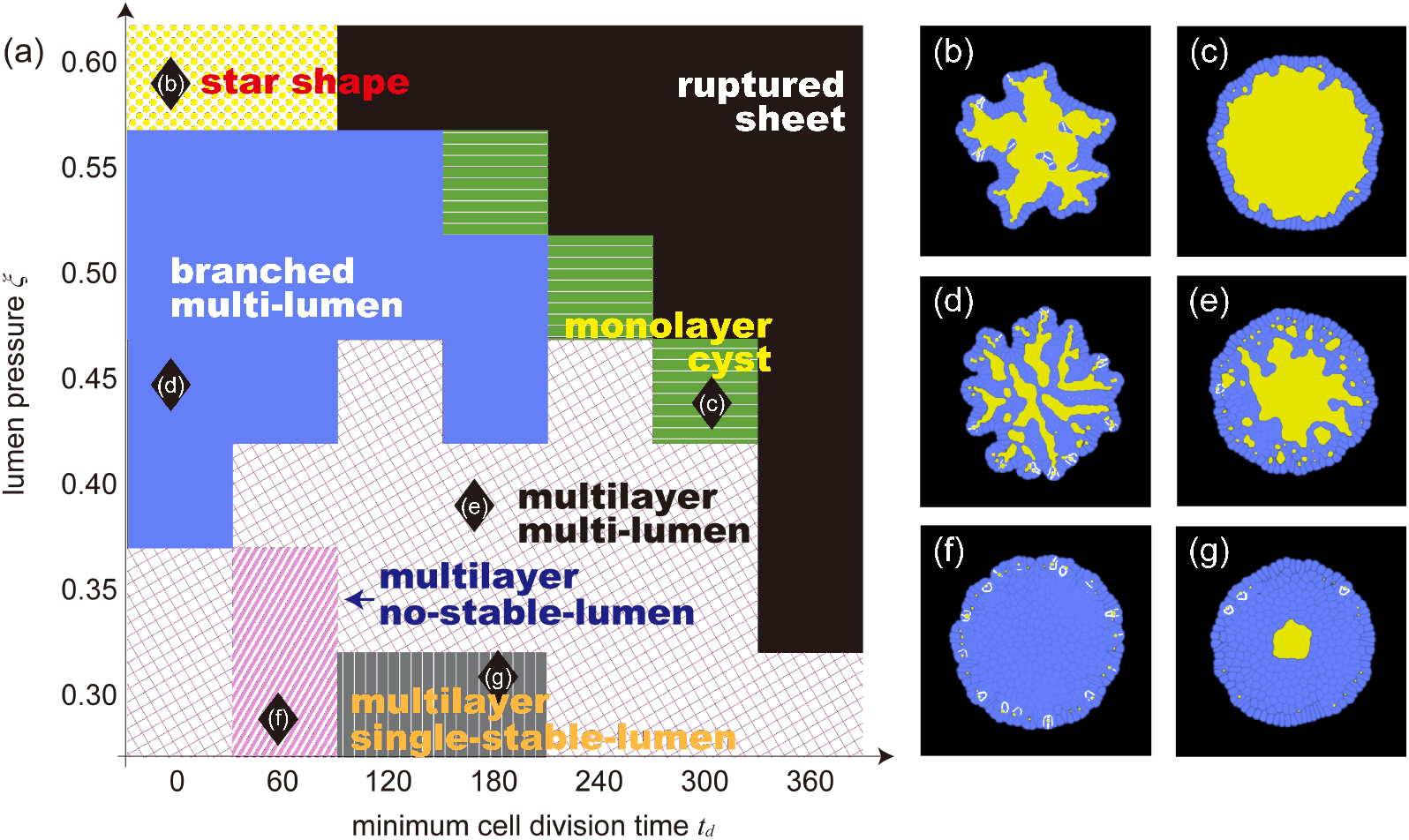
Morphologies resulting from the addition of noise to the volume condition for cell division. (a) A phase diagram of the organoid morphology and typical pattern of each phase with added noise to the volume condition for cell division. Each domain corresponds to: star shape (yellow dots), monolayer single-lumen (green horizontal stripe), branched multi-lumen (red solid), multilayer multi-lumen (magenta lattice), multilayer no-stable-lumen (blue tilted stripe), and multilayer single-stable-lumen (navy vertical stripe). The black diamond markers correspond to the parameter sets where the organoids of (b)-(g) emerge. The final states of (b) star shape organoid formed when (*ξ, t*_*d*_) = (0.60, 0), (c) monolayer single-lumen organoid formed when (*ξ, t*_*d*_) = (0.45, 300), (d) branched multilayer organoid formed when (*ξ, t*_*d*_) = (0.45, 0), (e) multilayer multi-lumen organoid formed when (*ξ, t*_*d*_) = (0.40, 180), (f) multilayer no-stable-lumen organoid formed when (*ξ, t*_*d*_) = (0.30, 60), (g) multilayer single-stable-lumen organoid formed when (*ξ, t*_*d*_) = (0.30, 180).

In addition to observed morphological differences, it is important to consider the probability of the cell layer rupturing. For the simulations with added noise at *t*_*d*_ = 300, we observed rupturing events in the monolayer regions. Specifically, out of ten simulations, there were two instances of rupturing at *ξ* = 0.35, five instances at *ξ* = 0.4, seven instances at *ξ* = 0.45, and nine instances at *ξ* = 0.5. It is worth noting that although the patterns without rupturing at *ξ* = 0.35 were eventually classified as multilayer in a later phase of the simulation, the rupturing occurred while the organoid was still a monolayer during the initial phase.

Notably, the branched multi-lumen morphology closely resembled that of the pancreatic organoid as shown in Fig 1. Conversely, the multilayer no-stable-lumen and multilayer single-stable-lumen morphologies remained unchanged. In summary, the morphology with added noise was more similar to the morphology observed in *in vitro* experiments than the morphology of those without added noise.

### Mechanism to maintain a monolayer with a monolayer single-lumen

As discussed in the Materials and Methods section, our model does not have a specific feedback mechanism implemented to maintain a monolayer. However, we discovered that a monolayer often emerged within a considerable parameter space, particularly when the *ξ* and *t*_*d*_ were large, as shown in Fig. 2(a) and 7(a). It is important to note that this monolayer remains present throughout the growth process, rather than only in the final simulation phase. The question arises about why this monolayer state emerges in the broad range of parameter space.

To better understand this mechanism, we examined how the lumen perimeter *ℓ*_*l*_ changes over time in relation to the number of cells in the organoid *N* . In Fig. 8(a), the red line represents the time evolution of *ℓ*_*l*_ at (*ξ, t*_*d*_) = (0.33, 300). It shows good agreement with the plot of *aN* (red line), which assumed a typical apical surface area of 1.6 arb. unit of length. This indicates that the rate of cell division is balanced with the rate of lumen growth, which results in the maintenance of a monolayer and enclosure of the lumen. If the rate of cell division is higher than the rate of lumen growth, a multilayer state may form. Conversely, if the rate of cell division is lower than the rate of lumen growth, the cell layer may rupture.

**Fig 8.**
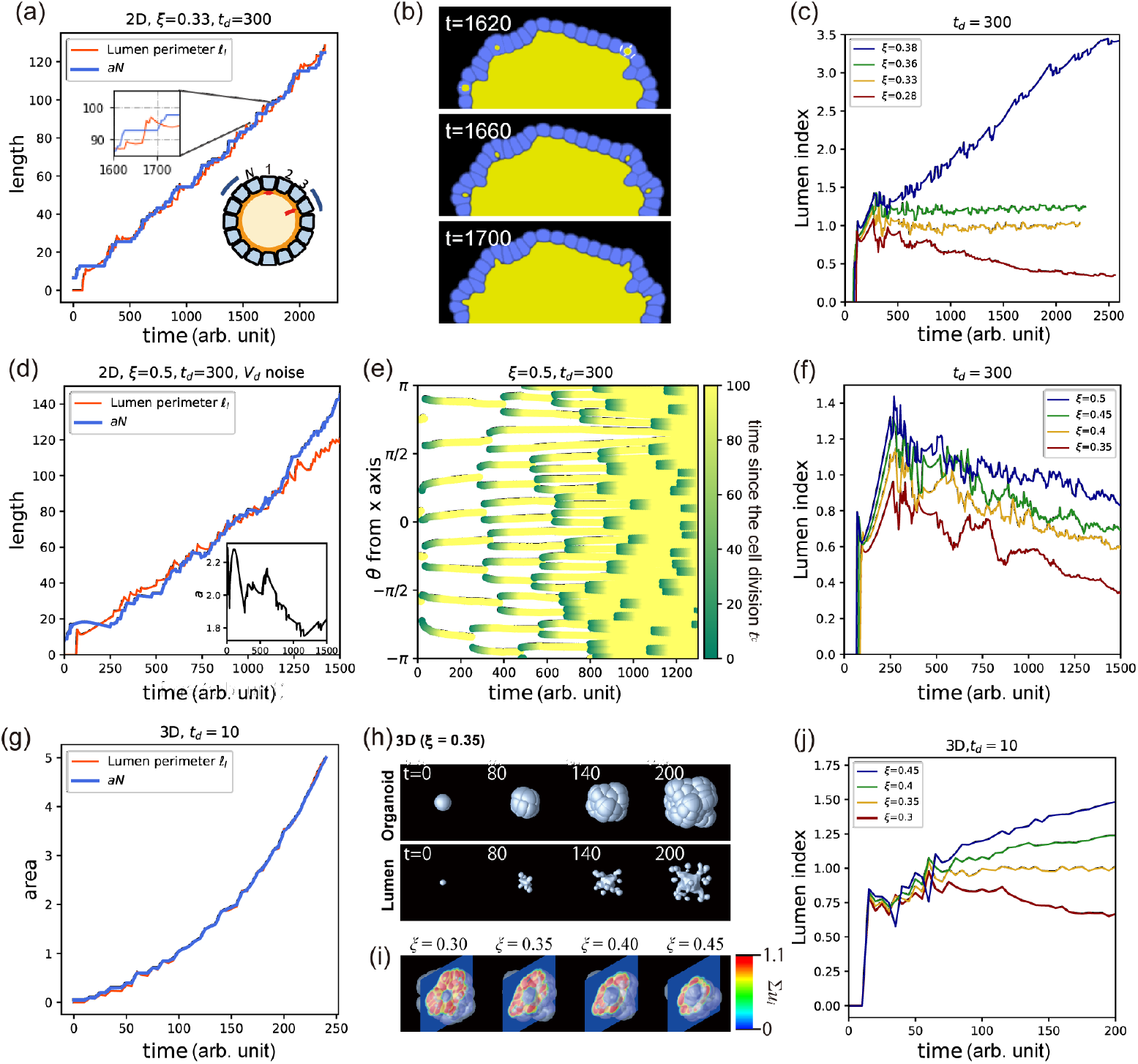
Process of maintaining monolayers; (a) Lumen perimeter *ℓ*_*l*_ and the number of cells *N* were plotted as a function of time for the 2D simulation with *ξ* = 0.33 and *t*_*d*_ = 300. Note that *N* (*t*) has been multiplied by a constant factor of 1.6 to align with the curve of *ℓ*_*l*_(*t*) on the right vertical axis. Inset: An enlarged view of the two curves. Inset: *N* (*t*) shows a step-wise increase while *ℓ*_*l*_(*t*) exhibits a rapid increase followed by a decay. (b) Micro-lumens were generated just after cell division and merged into a central lumen at *ξ* = 0.33 and *t*_*d*_ = 300. (c) Lumen index, defined by *a* = 1.6 and *t*_*d*_ = 300, was plotted for various values of *ξ*. (d) The lumen perimeter *ℓ*_*l*_ and the number of cells multiplied by the short axis of a cell *aN* were plotted as a function of time for the 2D simulation with added noise to the volume condition and with *ξ* = 0.45 and *t*_*d*_ = 300. Inset: Variation of the typical cell width *a* as a function of time. (e) A kymograph of the cell division events was plotted as a function of the angle coordinate. The color indicates the elapsed time since the cell was generated through a cell division. (f) The lumen index, defined by *t*_*d*_ = 300 and with added noise to the volume condition, was plotted for various values of *ξ*. (g) The lumen surface, defined by Eq.(4), *S*_*l*_, and the number of cells *N* were plotted as a function of time for the 3D simulation at (*ξ, t*_*d*_) = (0.35, 10). Note that *a* = 0.05. (h) The evolution of the organoid shape is shown for the 3D simulation with (*ξ, t*_*d*_) = (0.35, 10). (i) The lumen index, *S*_*l*_*/aN*, was plotted for the 3D simulation with various lumen pressure values *ξ* = 0.30, 0.35, 0.40, and 0.45 and *t*_*d*_ = 10.

As shown in the inset of Fig. 8(a), *ℓ*_*l*_ increases sharply when *N* is in the plateau state, but decreases as cell division occurs and the *N* increases. This interplay between the two curves is due to the volume exclusion interactions between the cells and lumen. When there is a large number of cells in the layer, they push against each other, thus preventing cell volume growth beyond a minimum volume required for cell division (*V*_*d*_). However, when the lumen enclosed by the cell layer grows, it stretches the perimeter of the cell layer, thereby creating more space for cells to grow outward. When a cell reaches the minimum *V*_*d*_ required for division, it divides, and its daughter cells then rapidly increase in volume. This growth pushes the micro-lumen generated when the cell is divided backward until it expands again as the growth rate of the cell volume decreases, and the micro-lumen merges into the larger lumen, as shown in Fig.8(b).

To examine the time evolution of the relation between the lumen perimeter and number of cells, we introduced a new index called the lumen index, which is defined as

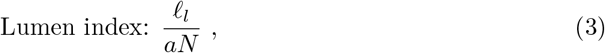

where *a* is a typical apical surface width of a cell in two dimensions. The apical surface is defined as the surface of a cell facing inward toward to the lumen area. The time derivatives of the lumen index reveal the following important dynamics: a negative coefficient indicates narrowing of the cell’s apical surface area or an increase in the number of cell layers, while a positive coefficient suggests widening of cell width, monolayer rupture, or merging of the micro-lumens.

In Figure 8(c), we observed the lumen index for various lumen pressures, with an apical surface area of 1.6 arb. units of length. The organoids at *ξ* = 0.32 and 0.36 exhibited a monolayer single-lumen state, and their lumen index remained close to one. The lumen index at *ξ* = 0.28 decreased with time, and the number of cell layers increased. The lumen index at *ξ* = 0.38 increased with time, while the cell layer ruptured at around *t* = 500.

A similar correlation between the lumen perimeter and number of cells was observed when noise was added to the minimum volume condition for cell division. Figure 8(d) shows the time evolution of *ℓ*_*l*_ and *aN* at (*ξ, t*_*d*_) = (0.5, 300), where *a* was defined as the average short axis of each cell when fitted with an ellipse, over all the cells present on the organoid at that time. Similar to the previous case, *ℓ*_*l*_ and *N* increase with time at similar rates. A standing point is when cell division across uniformly on the cell sheet, as indicated by the angular coordinates of cells at each time point plotted in Fig. 8(e), where the colors indicate the time elapsed since the cell was generated. Therefore, the morphology became rounder compared with the similar conditions without noise.

Figure 8(f) presents the lumen index at *t*_*d*_ = 300 for the different values of *ξ*=0.35, 0.4, 0.45, and 0.5. A monolayer single-lumen was stably formed at *ξ* = 0.5 and exhibited a lumen index of around 1.0, while the others switched from a monolayer to multilayer. Those at *ξ*=0.35, 0.4, and 0.45 remain monolayered until approximately 800, 1000, and 1100 arb. units of time, respectively. After 1000 arb. units of time, the number of cells that did not face the lumen increased, leading to a decrease in the lumen index.

The monolayer stabilization mechanism has been widely discussed in the literature, with the stretch-induced cell proliferation being a proposed mechanism [48–51]. The concept is that, if the stretching of tissue increases the rate of cell proliferation, the monolayer will be robustly maintained, and this process may be controlled through mechanobiological feedback mechanisms. However, our simulations suggest that the minimum cell volume condition for cell division may have a similar effect through the direct mechanical interactions between the cells and lumen.

We further confirmed this mechanism in the 3D simulation of the phase field model with *t*_*d*_ = 10, as shown in Fig. 8(h). The parameters used for the 3D simulations were almost the same as those in the 2D simulations, except for *V*_*d*_ = 3.88 and *V*_*target*_ = 4. In the 3D simulation, we plotted the time evolution of the number of cells *N* and lumen surface area defined by the equation:

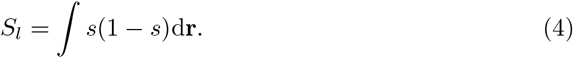

As shown in Fig. 8(g), the two curves overlap nearly perfectly as a function of time (*ξ* = 0.35). Figure 8(h) depicts the outer layer shapes of the organoid and lumen surface. Although numerous micro-lumens can be observed, only one stable lumen is present in the center. Figure 8(i) displays cross-sectional views of the organoids at t=200 arb. units of time for the different values of *ξ*. The organoid at *ξ* = 0.3 exhibits multiple layers, whereas those at *ξ* = 0.35 and 0.4 maintain a monolayer structure. The organoid at *ξ* = 0.45 is also a monolayer, but some cells appear to be stretched.

We also investigated the time evolution of the lumen index for the various values of *ξ* in 3D, as shown in Fig.8(j). Here we used *a* = 0.05. At *ξ* = 0.3, the lumen index decreased over time, and at this parameter, the organoid had a multilayer structure when the number of cells was 85, as shown in Fig.8(i). At *ξ* = 0.35 and 0.4, the lumen index remained around specific values. These results suggest that the organoids can maintain a monolayer structure, as shown in the cross-section of Fig.8(i), at least in the observed period. At *ξ* = 0.45, the lumen index increased over time, suggesting that the cell layer rupture is expected to occur at a specific time.

The lumen index is a useful parameter to observe the cell layer dynamics in experimental systems. As demonstrated in our simulation, morphological changes such as the timing of transitioning from a monolayer can be identified by tracking the time course of the lumen index. The value used to represent the cell width in the simulation, *a*, can be substituted with an appropriate measure in the experiment. For example, if the cross-sectional area of a typical cell in the system is known, then its square root can be used as a surrogate for cell width. Conversely, in a system where a monolayer is known to exist, changes in the cell membrane can be assessed by measuring the apical surface area of the cells using the *L/N* ratio. Furthermore, the time derivative of the lumen index allows for the observation of micro-lumen fusion. Analyzing the temporal relationship between this measurement and phenomena such as cell division provides valuable biological insights.

### Mechanism of forming a monolayer star-shape

This subsection discusses the mechanism for the formation of star-shaped organoids. As shown in Fig. 9(a), the star-shaped organoid at (*ξ, t*_*d*_) = (0.35, 300) without added noise in the *V*_*d*_, exhibited a cellular layer protruding outward. In addition, some cells extruded toward the lumen. In our simulations, this shape with broken rotational symmetry emerged spontaneously.

**Fig 9.**
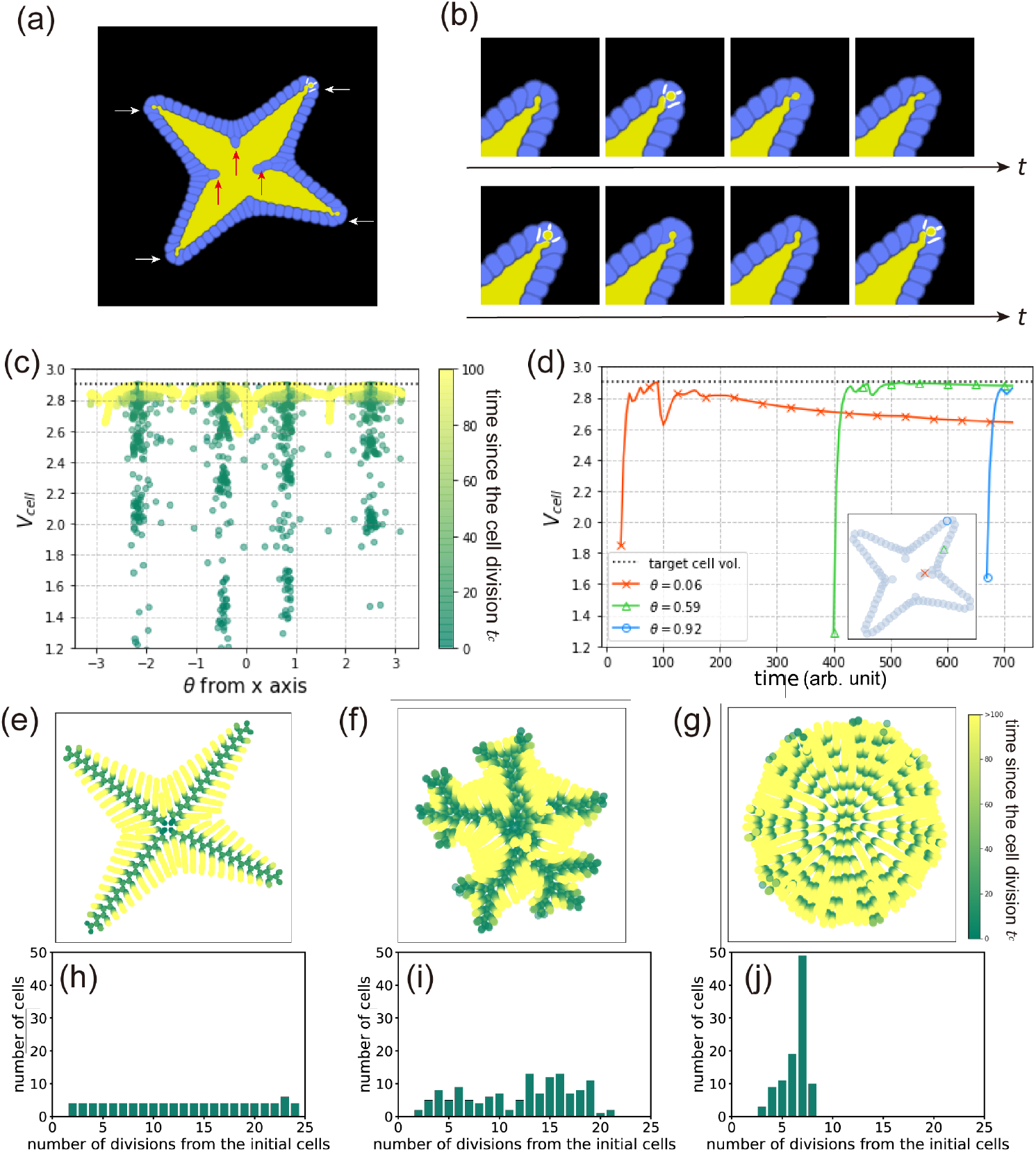
Mechanism for the formation of a star-shaped organoid. (a) Snapshot of the typical morphology of a star-shaped organoid formed at (*ξ, t*_*d*_) = (0.35, 0). Red arrows indicate cells extruding into the lumen, and white arrows show the branches of the organoid. (b) Time evolution of a branch tip. (c) Time-integrated distribution of cell volume of the cells for angular coordinates of the cells appearing in a star-shaped organoid at (*ξ, t*_*d*_) = (0.35, 0). The vertical lines represent the angles of the lumen branches. The horizontal line represents the volume of the cells required for cell division. (d) Relations between the cell volume and position of the representative cells in a star-shaped organoid at (*ξ, t*_*d*_) = (0.35, 0). The orange cross, green triangle, and blue circle markers represent typical cells in the center, on an arm, and at the tip of an arm, respectively. Markers in the inset represent the final positions of each cell; (e-g) Historical positions of each cell. All cells at the various times are overlaid in the same figure, and the color indicates the elapsed time from the generation of each cell at each time *t*_*cell*_. (h-j) Histograms of the frequencies of the number of divisions from the initial cells. Simulation conditions for panels (e-j) are as follows: (e) and (h) are without added noise to ‘*V*_*d*_’ and at (*ξ, t*_*d*_) = (0.35, 0); (f) and (i) are with added noise to ‘*V*_*d*_’ and at (*ξ, t*_*d*_) = (0.60, 0); and (g) and (j) are with noise to ‘*V*_*d*_’ and at (*ξ, t*_*d*_) = (0.45, 300).

Observing the growth at the tip, it appears that only the cells arranged at the tip could divide [Fig. 9(b)]. Cells along the elongated branches were compressed in the tangential direction, whereas the apical cells were not subjected to those compressive forces. Therefore, apical cells could grow.

To examine the relationship between the cell volume and position, we plotted the average cell volume at each angle position, as shown in Fig. 9(c). Various cell sizes were observed along the angles of the lumen branches, indicating that cells located along the branches surpassed the cell volume required for division and produce daughter cells. Conversely, in other areas, cells were unable to reach the required cell volume for division and remained undivided.

To investigate the reason why only cells at the tip of the branches had the ability to divide, we examined changes in the cell volume over time. Figure 9(d) shows changes in the cell volume and their position after the cells in the organoid were generated. Cells that were generated in the early stages of the simulation and extruded towards the lumen at the end of the simulation increased in area immediately after generation [orange crosses in Fig. 9(d)]. However, their volume did not increase to the extent required for cell division, *V*_*d*_, and subsequently decreased with time because of compression caused by the growth of subsequently-generated cells. Cells that remained in the cellular layer at the end of the simulation also increased in area to some extent after generation but did not surpass the cell volume required for cell division due to the compressive forces [green triangles in Fig. 9(d)]. In contrast, cells located at the tip of branches increased their volume to the required division volume condition immediately after generation and subsequently divided [blue circles in Fig. 9(d)].

Figure 9(e) shows the positions of each cell at the various times overlaid in the same figure for the parameters (*ξ, t*_*d*_) = (0.35, 300). The colors indicate the elapsed time from the generation of each cell. From the figure, it can be seen that new cells were only generated at the tips of the branches, as indicated by the concentration of dark green color at the central part of the branches. In contrast, cells that did not divide remained in other positions. These results suggest that cells spontaneously change their roles depending on their position, and only the cells at the branch tips have the ability to divide.

Furthermore, we analyzed the number of cell divisions from the initial cells in the final state at (*ξ, t*_*d*_) = (0.35, 300). Figure 9(h) shows a histogram of the number of divisions. For most bins, the number of cells is four, which is consistent with the number of branches. This finding suggests that only one of the two daughter cells generated through cell division is able to divide again, leading to a repetition of cell divisions that form each branch.

To compare the star-shaped organoid with other monolayer organoids, we examined the same plots for cases with added noise in the volume condition for cell division. At (*ξ, t*_*d*_) = (0.60, 0), where the organoid forms a more-branching star shape, the positions of cell divisions are consistent with the extension of the branches [Fig. 9(f)]. The histogram also indicates a state in which the cell lineages with division potential are limited [Fig. 9(i)]. In contrast, at (*ξ, t*_*d*_) = (0.45, 300), where the organoid forms a round monolayer with a single-lumen, cell divisions uniformly occur in the angular direction, particularly in the first half stage [Fig. 9(g)]. The histogram shows that most cells undergo approximately seven divisions since their initial cell stage and exhibit little variability. This is consistent with the division potential that is not coupled with the position [Fig. 9(j)].

Intestinal organoids exhibit a phenomenon similar to that of the spontaneous positioning and formation of tubular structures observed in stem cells. These organoids develop tubular branches from a single-lumen, with the cells that possess division ability located mainly at the tip of the tubular structure [31]. Theoretical studies have suggested that the coupling of cell division and curvature of a single cell layer contributes to the formation of the structures that are observed in the intestinal lining [52]. However, further research is required to determine whether organoids grow tubular branches from small outward protrusions because of positive feedback that arises from the division conditions, as demonstrated in this simulation.

## Discussion

In this study, we investigated the impact of mechanical elements on the organoid morphogenesis. We utilized a phase field model to simulate multicellular systems, where the cell-lumen interactions were mediated through various pressure conditions. We explored conditions with varying lumen pressure and minimum time required for cell division, and classified the morphology of the resulting organoids using these proposed indicators.

The growth rate of the observed organoids under the experimental conditions varied greatly depending on the system, making it challenging to directly compare the different systems. However, as demonstrated in this study, it is possible to compare them when measuring the number of cells, volume, and surface area of the organoid and lumen. This can be confirmed through scaling relations between the cell number versus the volume and surface area. For example, if the organoid is spherical with a radius of *R*, the volume and surface area of the organoid scale is *R*^3^ and *R*^2^, respectively. If an organoid is a monolayer, the number of cells should be proportional to surface area, *R*^2^, and if there is no lumen inside the organoid, then *N* ∝ *R*^3^. In other words, the relationship between the radius, volume, or surface area and the number of cells can be used to describe the organoid structure. The same principle can be applied to more complex organoid morphologies. For example, pancreatic organoids, which exhibit a monolayer organoid with branching, are known to show fractal dimension, and the relationship between the surface area and volume deviates from the aforementioned power-law relationship [22].

In the Results section, we went beyond simple scaling arguments and introduced novel indicators to characterize the morphology of the organoids grown under the various conditions. Specifically, we proposed the sphericity and lumen index, which capture the deviations from simple spherical shapes and highlight when lumen structure changes. Additionally, these indicators are also useful for experimental investigations as well. Our study thus offers a deeper understanding of how mechanical elements influence organoid morphogenesis and provides new tools to analyze and quantifying this complex process.

In addition, we began the simulations with four cells. While the quantitative impact of the initial cell number on the morphogenesis has not been thoroughly investigated in this model, we observed that starting with only one or two cells tended to increase the frequency of rupture events. This phenomenon can be attributed to the presence of small number of cells in the initial phase, making it challenging to effectively surround the lumen formed during the initial cell division. These preliminary results suggest the possibility of achieving specific morphologies by appropriately controlling the rate of lumen expansion during different formation stages, as also demonstrated in experimental studies [14].

Our phase field model was advantageous because it could easily incorporate the lumen pressure, making it possible to discuss the effect of lumen pressure as a variable of the organoid morphology. However, our current formulation cannot reproduce structures where the apical surface of the cells protrude toward the lumen, which is present in some organoids which show rosette-like structures. One way to address this issue is to introduce a model for molecules localized on the apical surface, which could be achieved by modifying the multicellular phase field model. Future research can focus on reproducing the curvature of the cell membrane. In addition, our model does not currently capture the network structure that is observed in the pancreas. To reproduce this network structure, it is necessary to incorporate a tube-shaped lumen formation mechanism into the model. This would require updates to the model, such as considering the reaction-diffusion processes [46]. Incorporating these additional elements could provide a more comprehensive understanding of the organoid morphogenesis and enable the simulation of the complex network-like structures that are observed in certain organs.

It is important to acknowledge that the current model represents organoids in a state where they can grow in an outward direction without any physical constraints. However, in real organoids or organs, growth is influenced by various factors, including the extracellular matrix (ECM). Mechanical interactions, which are the focus of this research, are also expected to be influenced by the external environment, such as the pressure exerted by the ECM on the cells as some experimental studies have demonstrated [53, 54]. Considering the pressure exerted by the ECM, it can be anticipated that the growth of cells in the outer shell of the organoid would tend to be more uniform, as some outer cells would experience resistance and they would be hindered from protruding outward. Exploring how these dynamics change in a model that incorporates the ECM would be a valuable avenue for future investigations.

In summary, we used a simple mathematical model to investigate the morphogenesis of organoids that were affected by mechanical factors. We introduced the minimum cell volume and minimum elapsed time as parameters and examined the organoid growth. The organoid morphologies were classified into seven distinct patterns based on the lumen occupancy, number, and sphericity. In addition, the dynamics of morphology can be monitored by lumen occupancy and lumen index. Those parameters are useful to characterize the morphologies observed in experiments.

## Materials and methods

We modeled the dynamics of cells and lumenal fluid using the phase field model, building on previous studies [37, 38]. To avoid excessive computation time, most of the simulations were performed in 2D, except for the simulations that are presented in Fig. 8(g-i).

### Time evolution

Similar to previous studies [37, 38], the existence of cells and lumens are described by the field values, *u*_*i*_ *∈* [0, 1]. Label *i* represents the index of a cell when *i ∈ {*1, 2, …, *M }* and a lumen when *i* = *L*, where *M* is the number of cells in the system. *u*_*i*_ = 1(= 0) designates the inside (outside) of the cell or lumen *i*. Common dynamics of the cells and lumens are described as follows:

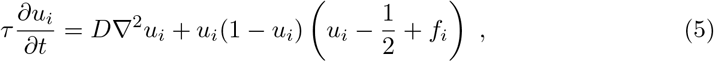

where the coefficient *τ* is a characteristic time for the field variables, diffusion coefficient *D* is proportional to the square of the interface thickness and *f*_*i*_ is a function that depends on whether the value represents a cell or the lumenal fluid. When *u*_*i*_ represents a cell, the function is described as follows:

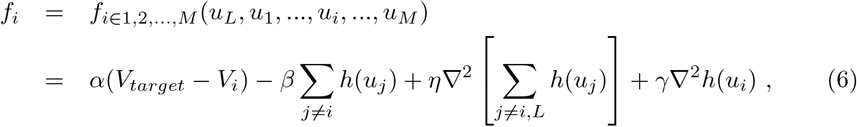

where the coefficients *α, β, γ, η*, and *V*_*target*_ are positive constants; *h*(*u*_*j*_) is a function that is defined as *h*(*x*) = *x*^2^(3− 2*x*) [37]; and *V*_*i*_ is the volume of the cell *Vi*. The volume of each cell, defined by *V*_*i*_ = *h*(*u*_*i*_)d*r*, increases towards the target volume *V*_*target*_ under the constraint of the excluded volume interaction (− *β Σ* _*j*=*i*_ *h*(*u*_*j*_) term) and adhesion interaction (3rd term on the right-hand side of Eq.(6)) with the other cells. When *i* = *L*, the function is described as follows:

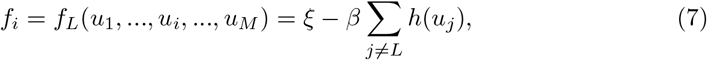

where the coefficient *ξ* represents the lumenal pressure, which is a positive constant. If multiple isolated lumen regions are in contact with each other, they are assumed to merge into one. See [37, 38] for additional details.

These equations were integrated numerically by discretizing them with d*x* in space and d*t* in time using the Euler scheme. All units used in this study are arbitrary units and not SI units such as m, s, or min. Rather, we use ‘arb. unit’ to represent the arbitrary units in this report. For ease of reference, we refer to the time of simulation as ‘time’ throughout this report.

### Cell division and micro-lumen creation

Cell division conditions can be represented using physical quantities, such as cell size and time since previous division. Different cell types have different division triggers, such as reaching a certain volume threshold or after a specific time period has elapsed since the last division [55]. In this study, we varied the time conditions as a control parameter.

We assumed that each cell was set to divide when both the volume and time conditions were met. The volume condition is formulated as *V*_*i*_ *> V*_*d*_, where *V*_*i*_ is the volume of the cell *i*, and *V*_*d*_ is the minimum cell volume required for cell division. In this study, we set the target cell volume *V*_*target*_ to 3.0 arb. unit of volume. For most of our simulations, *V*_*d*_ was fixed to 2.9 arb. unit of volume. We also implemented conditions where noise was added to the minimum volume condition, where *V*_*d*_ was set to follow a normal distribution with a mean of 2.85 and a standard deviation of 0.025. The time condition is formulated as *t*_*cell*_ *> t*_*d*_(1 + *ζ*). Here, *t*_*cell*_ is the time since the last cell division of the cell; *t*_*d*_ is the minimum time required for cell division, which is a positive constant, and *ζ* is white noise in the range of [− 0.1, 0.1]. The time when the volume condition is satisfied depends on the surrounding environment. Figure 10(a) presents an example of the time evolution of the cell volume. The presented cell is one of the two cells that were generated through the division of an isolated cell. The volume of the daughter cell increases rapidly after cell division and satisfies the volume condition at a certain elapsed time since the division. For instance, the elapsed time for a cell division from a single isolated cell was approximately 14 arb. unit of time. We refer to typical elapsed time as *t*_*cell*_(*V*_*d*_). When the cell time condition *t*_*d*_ is greater than *t*_*cell*_(*V*_*d*_), it divides at *t*_*d*_ [Fig. 10(b)]. Conversely, when *t*_*d*_ is less than *t*_*cell*_(*V*_*d*_), the cell divides at *t*_*cell*_(*V*_*d*_) [Fig. 10(c)]. If either of the conditions was not met, the cell is assumed to remain in its current state, without undergoing division or extinction, but retains the potential to change its volume and shape.

**Fig 10.**
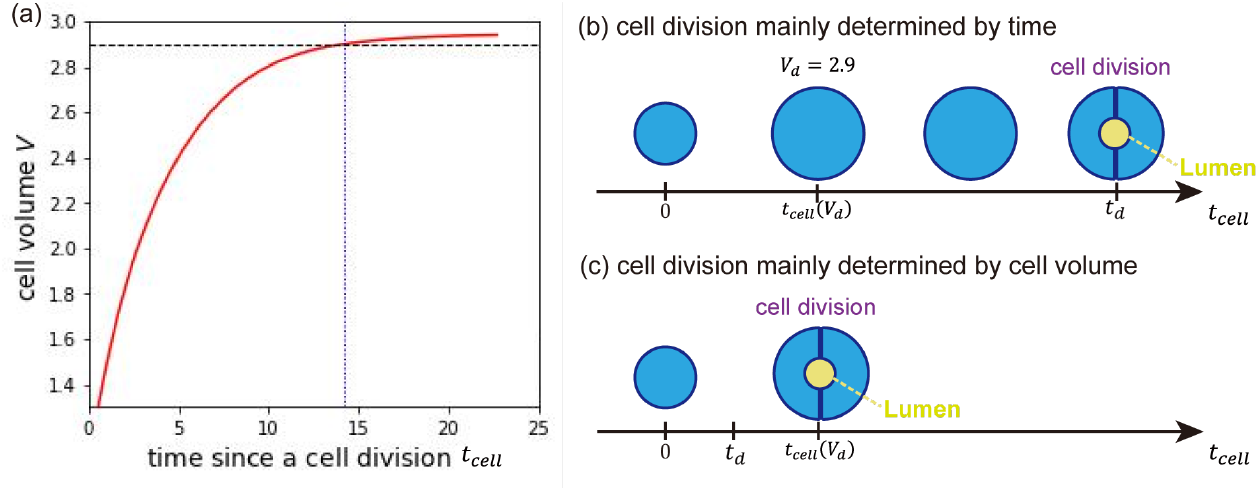
Figure illustrating the conditions that need to be met for cell division to occur in the model used for this study. (a) Time evolution of the cell volume of a daughter cell that was generated through cell division. The horizontal dashed line represents *V*_*d*_ = 2.9 arb. unit of volume, which is the minimum volume required for cell division to occur. The vertical dotted line indicates the time when the cell satisfies the volume condition for cell division. (b) When the time condition is dominant, the cell divides immediately after the time condition is satisfied. (c) When the volume condition is dominant, the cell divides immediately after the volume condition is satisfied.

The cell immediately divides when both time and volume conditions are satisfied. As per a previous study [38], the division plane is determined by the position of the two spindle poles. The position of the poles, represented by the position vectors ***r***_1_ = (*x*_1_, *y*_1_) and ***r***_2_ = (*x*_2_, *y*_2_), are determined by the steady state of the following equations;

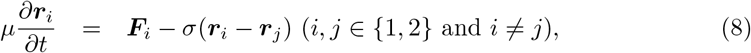

where *μ* and *σ* are positive constants. The force applied to the poles can be expressed as:

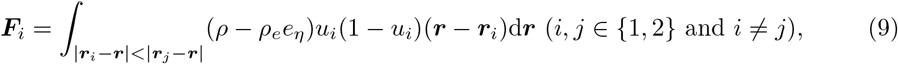

where *ρ* and *ρ*_*e*_ are positive constants, and *e*_*η*_ is a function given as

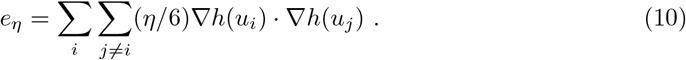

This force enables the division plane to be parallel to the cell-cell contact plane if the cell attaches to other cells. See [38] for additional details.

A spherical micro-lumen is created at the center of the division plane with a constant size of *V*_*L*,*ini*_ = 0.785 arb. unit of volume simultaneously with the determination of the division plane. It is assumed that the lumen does not exist until cell division occurs. As mentioned above, the generated lumens evolve over time according to the Eqs. (5) and (7). The values of parameters utilized in this investigation are shown in Table 2.

**Table 2.** Parameter values employed in this research (arb. unit). See [37, 38] for additional information.

| parameter | $D$ | $\tau$ | $\alpha$ | $\beta$ | $\gamma$ | $\eta$ | $\epsilon$ | $\sigma$ | $\mu$ | $\rho_0$ | $\rho_e$ |
| --- | --- | --- | --- | --- | --- | --- | --- | --- | --- | --- | --- |
| values | 0.001 | 1 | 1 | 1 | 1 | 0.008 | 0.1 | 0.001 | 10 | 0.01 | 1 |

## Acknowledgments

We would like to express our deepest gratitude to all individuals who contributed to the success of this research. We are also grateful to Markus Mukenhirn, Yara Alcheikh, Tzer Han Tan, Allison Lewis, Irene Seijo Barandiaran, Siham Yennek, Phil Seymour, Heike Petzold, and Jacques Prost for their insightful discussions and valuable input on the mechanism of forming a star shape monolayer. This work was initiated and supported by a Research Grant of the Human Frontier Science Program (GP0050/2018) to DR, AH, AGB, and MS. This work of the Interdisciplinary Thematic Institute IMCBio, as part of the ITI 2021-2028 program of the University of Strasbourg, CNRS and Inserm, was supported by IdEx Unistra (ANR-10-IDEX-0002), and by the SFRI-STRAT’US project (ANR 20-SFRI-0012) and EUR IMCBio (ANR-17-EURE-0023) under the framework of the French Investments for the Future Program to DR. This work was supported by the Research Fund for International Scientists of NFSC (12250710131) to MS and the Seed Fund of Mechanobiology Institute to TH, and JSPS KAKENHI (23K03345) to MN.

## Author Contribution

Conceptualization: Anne Grapin-Botton, Alf Honigmann, Daniel Riveline, Masaki Sano Formal Analysis: Sakurako Tanida

Funding Acquisition: Anne Grapin-Botton, Alf Honigmann, Daniel Riveline, Masaki Sano

Investigation: Sakurako Tanida, Makiko Nonomura Methodology: Makiko Nonomura, Tetsuya Hiraiwa, Masaki Sano Project Administration: Daniel Riveline, Masaki Sano

Software: Sakurako Tanida

Supervision: Daniel Riveline, Masaki Sano Validation: Kana Fuji

Visualization: Sakurako Tanida, Linjie Lu, Tristan Guyomar, Byung Ho Lee, Makiko Nonomura

Writing -Original Draft Preparation: Sakurako Tanida, Tetsuya Hiraiwa, Masaki Sano Writing -Review & Editing: Anne Grapin-Botton, Alf Honigmann, Byung Ho Lee, Daniel Riveline, Kana Fuji, Linjie Lu, Makiko Nonomura, Tristan Guyomar

